# Neural Responses in Dorsal Prefrontal Cortex Reflect Proactive Interference during an Auditory Reversal Task

**DOI:** 10.1101/354936

**Authors:** Nikolas A. Francis, Susanne Radtke-Schuller, Jonathan B. Fritz, Shihab A. Shamma

## Abstract

Task-related plasticity in the brain is triggered by changes in the behavioral meaning of sounds. We investigated plasticity in ferret dorsolateral frontal cortex (dlFC) during an auditory reversal task to study the neural correlates of proactive interference, i.e., perseveration of previously learned behavioral meanings that are no longer task-appropriate. Although the animals learned the task, target recognition decreased after reversals, indicating proactive interference. Frontal cortex responsiveness was consistent with previous findings that dlFC encodes the behavioral meaning of sounds. However, the neural responses observed here were more complex. For example, target responses were strongly enhanced, while responses to non-target tones and noises were weakly enhanced and strongly suppressed, respectively. Moreover, dlFC responsiveness reflected the proactive interference observed in behavior: target responses decreased after reversals, most significantly during incorrect behavioral responses. These findings suggest that the weak representation of behavioral meaning in dlFC may be a neural correlate of proactive interference.

**Significance Statement:** Neural activity in prefrontal cortex (PFC) is believed to enable cognitive flexibility during sensory-guided behavior. Since PFC encodes the behavioral meaning of sensory events, we hypothesized that weak representation of behavioral meaning in PFC may limit cognitive flexibility. To test this hypothesis, we recorded neural activity in ferret PFC, while ferrets performed an auditory reversal task in which the behavioral meanings of sounds were reversed during experiments. The reversal task enabled us study PFC responses during proactive interference, i.e. perseveration of previously learned behavioral meanings that are no longer task-appropriate. We found that task performance errors increased after reversals while PFC representation of behavioral meaning diminished. Our findings suggest that proactive interference may occur when PFC forms weak sensory-cognitive associations.

## Introduction

An important role of prefrontal cortex (PFC) is to flexibly encode sensory-cognitive associations during goal-oriented behavior. Single-units in PFC rapidly adapt their responses to encode new behavioral meanings of sensory events, which is believed to facilitate successful behavior when task demands change (Asaad et al., 1998; Donahue and Lee, 2015; Durstewitz et al., 2010; Fritz et al., 2010; Kusunoki et al., 2010; Lee et al., 2009; Mian et al., 2014; Rigotti et al., 2013; Rodgers and DeWeese, 2014; Stokes et al., 2013). For example, individual units may become selective for task-related information that did not previously drive responses (Asaad et al., 1998; Donahue and Lee, 2015; Durstewitz et al., 2010; Fritz et al., 2010; Karlsson et al., 2012; Stokes et al., 2013; Watanabe et al., 1992). In addition, opponent cell populations that encode contrastive task-related information may emerge during behavior (Cromer et al., 2010; Kusunoki et al., 2010; Rodgers and DeWeese, 2014; Roy et al., 2010; Stokes et al., 2013; Zhou et al., 2016).

However, it remains unclear how modulation of PFC responsiveness correlates with complex behavior involving reversals of stimulus meaning that can introduce proactive interference, i.e., perseveration of previously learned behavioral meanings that are no longer task-appropriate. It is possible that proactive interference is partly due to the inability of PFC to fully update sensory-cognitive associations during task performance. Human functional neuroimaging studies have shown that PFC participates in resolving proactive interference during unexpected changes in semantic categories (Dolan and Fletcher, 1997), conflicting stimulus associations (Badre et al., 2005; Jonides et al., 1998; Nee et al., 2007; Wolf et al., 2010), or when there is an incongruence of task rules (Hyafil et al., 2009).

Based on our previous work demonstrating that ferret dorsolateral frontal cortex (dlFC) encodes the behavioral meaning of sounds (Fritz et al., 2010), we hypothesized that the inability of dlFC to rapidly track changing behavioral meanings might be a neural correlate of proactive interference.

We explored the relationship between behavior and dlFC responses by training ferrets on an auditory reversal task. Reversal tasks are a powerful tool for studying the neural basis of proactive interference because the range of motor responses and the stimuli are fixed; yet the behavioral meaning of stimuli changes across time during task reversal. Thus, reversal tasks control for motor- and stimulus-specific neural responsiveness, while emphasizing a conflict of opposing sensory-cognitive associations. In our go/no-go reversal task, the behavioral meaning (i.e., “go” vs. “no-go”) of high and low frequency tones was reversed across blocks of trials. We found that the animals’ overall motivation to perform the reversal task, and their ability to detect both low and high frequency tones was similar before and after reversals, as evidenced by similar task performance for both low and high go-signaling tones throughout the task. However, no-go target recognition was significantly worse after reversals. Based on this behavioral evidence of proactive interference, we hypothesized that dlFC responses to targets might also be weaker after reversals.

To study the relationship between proactive interference and neural responses in dlFC, we recorded spiking activity while the ferrets performed the reversal task. We found that neural responses in ferret dlFC were more strongly time-locked to sensory events rather than to motor events, and were more selective for the behavioral meaning of tones than for acoustic features. Similar to the animals’ impaired target recognition after reversals, dlFC responses to targets also decreased after reversals, especially during incorrect behavioral responses. These findings suggest that the neural activity in dlFC that encodes the behavioral meanings of sounds before reversals, may also contribute to proactive interference after reversals.

## Materials and Methods

Neural activity was recorded in dorsolateral frontal cortex (dlFC) of 2 awake, behaving ferrets. All experimental procedures conformed to standards specified by the National Institutes of Health and were approved by the University of Maryland Animal Care and Use Committee.

Animals were trained to discriminate between high and low frequency pure-tones for water reward (Figure 1). The animals were initially trained in sound-attenuated testing booths, where they could move freely in a small arena. Once they reached criterion on the task, they were implanted with a head-post and trained to perform the same task in a head-fixed preparation to enable stable neurophysiological recordings during behavior. Behavior and stimulus presentation were controlled by custom software written in MATLAB (MathWorks).

**Figure 1.**
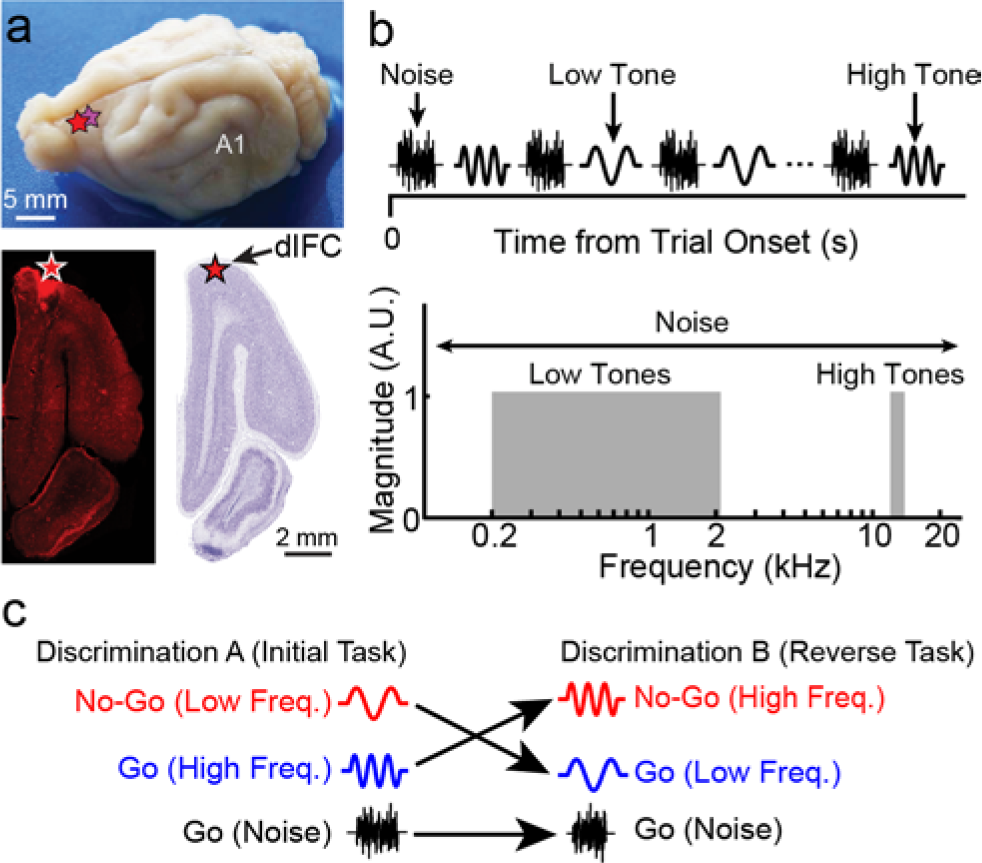
Dorsolateral frontal cortex (dlFC) recording sites and stimulus paradigm. **a.** The top panel shows a whole ferret brain, primary auditory cortex (A1) and the location of dlFC recording sites (in both animals) labeled with stars. Recording sites were verified as being in dlFC by examination of a fluorescent dye and/or an electrolytic lesion placed at the recording site of each animal. The lesions and region of high fluorescence were localized in left hemisphere coronal slices (lower left panel), then mapped onto an atlas of the ferret brain (lower right panel). The recording sites were localized to the dlFC, within the proreal gyrus. **b.** Each trial during the task consisted of either a noise or a tone. Noises and tones were alternated, and each tone presentation was randomly selected to come from either a low (200-2000 Hz) or high (13200-15000 Hz) frequency region, shown in the lower panel. The noise was broad-band and covered the range of tones (150-38400 Hz). **c.** During the reversal task, the behavioral meaning of the tones (“go” vs. “no-go”) were reversed across blocks of trials, while the behavioral meaning of the noise (“go”) was fixed throughout the task.

### Acoustic stimuli

Pure-tones were sine waves (5 ms onset and offset ramps; 1.0 s duration), with frequencies randomly selected from either a low (200-2000 Hz) or high (13200-15000 Hz) frequency band on each trial. Each pure-tone presentation, in a trial sequence of acoustic stimuli, was alternated with broad-band noise (150-38400 Hz). The silent inter-stimulus interval (between all stimuli) was 0.4 s. Noise durations were randomized in duration between 1.5-2.25 seconds. Sound levels were the same for all stimuli within an experiment, and ranged between 60 to 80 dB SPL across experiments. All sounds were synthesized using an 80 kHz sampling rate, and were presented through a free-field speaker that was equalized to achieve a flat gain from 200 Hz to 40 kHz.

### Auditory reversal task

Ferrets were trained on a conditioned avoidance “go/no-go” pure-tone frequency discrimination task that included reversals of behavioral meaning for pure-tones. During each trial, the animal licked a waterspout with a continuous flow of water while sounds were presented. During the Initial task, the low tones were assigned to be targets signaling avoidance, i.e. “no-go”—and the animals learned to stop licking the waterspout within 600 ms of target onset to avoid a mild shock. Conversely, the noise and high tones signaled positive reward during the Initial task, and evoked “go” behavior - approaching the waterspout to drink. Each behavior session began with a block of “Priming A” trials, during which only go-signaling sounds (high tones and noise) were presented. Following Priming block A, there was a block of go/no-go trials, the Initial task, during which the noise, target (low tones) and non-target (high tones) stimuli were presented. After completion of the Initial task, the animals received a second “Priming B” block of trials, during which a different set of go-signaling sounds (low tones and noise) were presented that prepared the animal for the upcoming task reversal. Following Priming B, the animals performed the “Reverse” task, in which the behavioral meanings (i.e. go vs. no-go) of the high and low tones were reversed (high tones were now target, and low tones were nontarget). We were not able to test responses in additional reversals, because the ferrets were no longer willing to perform additional pure-tone trials after the Reverse task. Hence, our main analysis focused on single-unit data from the two go/no-go blocks of trials (the Initial and Reverse tasks), during which the full stimulus set was presented, and there was a full reversal of pure-tone behavioral meanings. The statistical significance of behavioral data was determined with t-tests.

### Neurophysiology

Under anesthesia, each animal was surgically implanted in sterile conditions with a stainless steel head-post to allow for stable recording. Following recovering from surgery, a small craniotomy (1-2 mm diameter) was performed over dlFC. Neurophysiological recordings were conducted over a total of 14 behavioral sessions (7 for each animal). We used high impedance (1-7 MΩ) tungsten electrodes (FHC) for the neurophysiological recordings. Each recording session used four independently moveable electrodes (Alpha-Omega), separated by 500 μm in a 2 × 2 grid. Electrodes were advanced until spontaneous spiking was visible in voltage traces on the majority of electrodes. Data acquisition was controlled using custom MATLAB software. Single-unit responses were sorted offline using custom MATLAB software. Typically, it was possible to extract 1-3 separate and distinct stable waveforms (single-units) from each electrode.

### Data analysis

#### Single-unit response-type classification

Classification of responses as being either primarily auditory-, motor- or non-auditory-motor was done using a stepwise linear regression of spiking against auditory and motor (licking) events, binned at 20 bins/s. A unit was classified as primarily “auditory” only if the full regression model predicted spiking activity significantly better than the model based only on motor events (p < 0.05, t test). We focused our main analysis on auditory units—the population exhibiting the most consistent responses time-locked to sensory events.

#### Post-stimulus time histograms

Single-unit responses to task stimuli were measured by computing the post-stimulus time histograms (PSTH) of spiking. For consistent analysis, only the first 1.0 s of responses were considered, even when longer stimuli (for broad-band noise) were used. Responses were binned at 20 bins/s. A normalized response was computed for each unit by subtracting the baseline firing-rate from the PSTH, and dividing by the maximum absolute modulation from the spontaneous baseline.

#### General linear model

Neural responses in dlFC of awake behaving animals may be driven by many possible task-related variables. To contrast the influence of stimulus frequency, stimulus behavioral meaning, and the animals’ behavioral decisions in determining modulation from baseline spiking activity, we used a general linear model (GLM) of spiking, binned at 20 bins/s. Single-unit response values were taken from the average neural responses during each trial’s decision time-window, which was the 600 ms interval beginning with the onset of a sound and ending with the start of the potential punishment time-window. The model was solved in a least-squares sense (Haase, 2011) for each single unit, and the statistical significance of model parameters for each explanatory variable was determined using a bootstrapped t-test, with 10,000 resampling iterations.

### Verification of recording sites

Recording sites were documented by stereotaxic measurements that were referenced to fluorescent Di-I injections and lesions placed in the left frontal cortex (Figure 1A) as markers. Following histological processing, the markers were located in frontal brain sections and mapped onto an atlas of the ferret brain (Radtke-Schuller et al., unpublished). In both animals the recording areas were located within dlFC. In one animal, recording sites spanned 2100 μm to 3600 μm from the frontal hemisphere pole. The electrolytic lesion and collocated Di-I fluorescence were found 2400 μm from the frontal pole. In the other animal, the visible lesions spanned 2700 μm to 4200 μm from the frontal pole. The caudal-most region of the recordings in this animal may have been within a transitional zone between dlFC and premotor cortex however, premotor responses have previously been shown to be very similar to responses in dorsolateral PFC and to carry task-related auditory information (Lemus et al., 2009; Fritz et al., 2010). Nevertheless, in order to eliminate the possibility of motor-related spiking in our results, the stepwise regression model removed single-units that were classified as having primarily motor-driven responses (5/113 units).

## Results

We studied the relationship between proactive interference and single-unit responses in the dorsolateral frontal cortex (dlFC) of two ferrets trained to perform an auditory reversal task (Figure 1). We recorded from a total of 113 single-units in dlFC during task performance. Seven recordings were done in each behaving ferret. A stepwise linear regression model (see Materials and Methods) was used to classify units as being either primarily auditory (66/113, 58%), motor (5/113, ~4%) or non-auditory-motor selective (42/113, 37%). Since we were interested in how auditory information is represented in dlFC, and did not want to bias population statistics by responses that were correlated with non-auditory events, we focused our analysis on auditory units. In general, non-auditory unit responses were poorly timed-locked to acoustic events, and the overall magnitude of responses was <25% of the auditory units.

During task performance, over two thirds of the auditory units (n=45/66, 68%) responded in an excitatory (enhanced) manner to the target tones. We found only a few units with suppressed target responses (n=4/66, 6%), and the remaining population had neither consistent responses for a given stimulus, nor significant responses during decision periods of the task. Thus, further analysis was constrained to the 45 auditory units in dlFC with excitatory target responses (n=31 in ferret 1; n=14 in ferret 2).

### Proactive interference during reversal task performance

Animals were trained on a pure-tone frequency discrimination task that included reversals of behavioral meaning for pure-tones (Figures 1b and c). During the Initial task, low frequency tones signaled avoidance of waterspout licking (no-go), while the noise and high tones signaled approach (go). In contrast, during the Reverse task (following the Initial task), the behavioral meanings of the high and low tones were reversed, so that the animal now approached the waterspout for reward during low tones and noise, and avoided the waterspout during high tones. Thus, noise was always an approach signal, whereas the behavioral meaning of tones changed from the Initial task to the Reverse task.

Task performance was assessed by measuring licking behavior during the presentation of sounds. The top panel of Figure 2a shows the time-course of the likelihood of licking during each trial, for each stimulus. The bottom panel of Figure 2b shows task performance rates that summarize licking behavior during the decision time-window, which was the 600 ms interval beginning with the onset of a sound and ending with the start of the potential punishment time-window. We defined the “safe rate” as the likelihood of licking during a noise stimulus, which was expected to be high since the noise was always an approach signal. The safe rate therefore was a measure of the animal’s overall motivation to lick. The average safe rates during the Initial and Reverse tasks were not significantly different (99.9% ± 0.2% and 99.5% ± 1%, respectively; p=0.18, t-test), indicating that the animals were highly motivated to lick the waterspout throughout both tasks. The false alarm rates (i.e. the cessation of licking upon detecting a non-target) for the Initial and Reverse task were also comparable (38.4% ± 21.2% and 34.0% ± 16.4%; p=0.38, t-test). However, the hit rate (i.e. the cessation of licking upon detecting a target) decreased during the Reverse task, compared to the Initial task (83.6% ± 5.6% and. 76.9% ± 4.8%, respectively; p=0.009, t-test).

**Figure 2.**
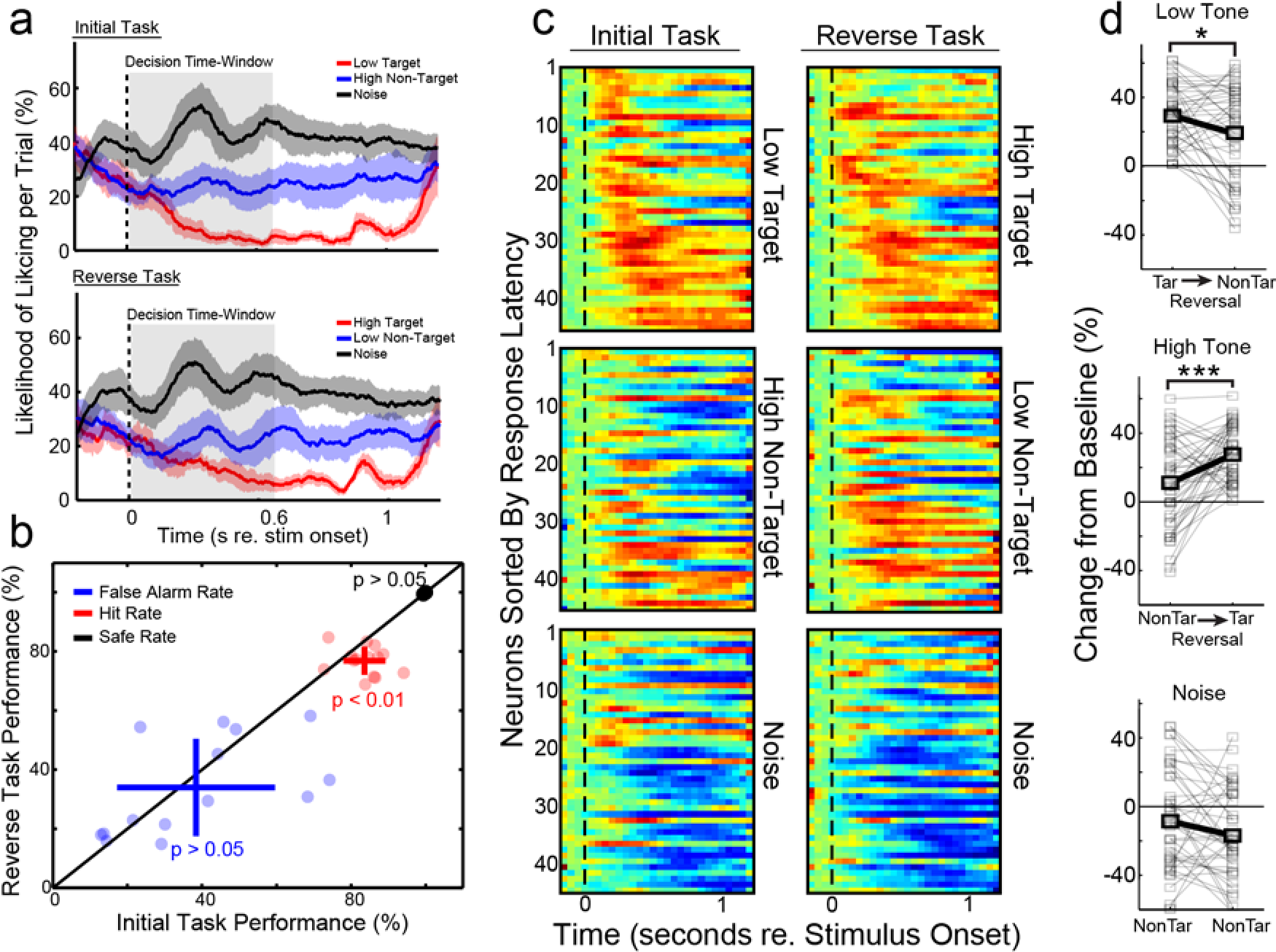
Reversal task performance and adaptive plasticity in dlFC. **a.** The top and middle panels show the likelihood of licking per trial for noise (black line), targets (red line) and non-targets (blue line). The middle panel shows the licking during the Reverse task, in which the behavioral meanings of the low and high tones were reversed compared to the Initial task (top panel). The vertical dotted lines show the stimulus onsets. Shade around lines shows 2 standard deviations of the mean (σ). The grey rectangles show the behavioral decision time-window, which was the 600 ms after the stimulus onset, until the start of the potential punishment time-window. **b.** The values for false alarms, hits and safe rates are shown in blue, red and black, respectively. For each statistic, each dot shows the result for a single experiment. Crosses show the performance means and standard deviations. The safe rate data were very consistent across experiments, and so they appear as a dot. p-values > 0.05 indicate insignificant differences between the Initial and Reverse tasks. Only the hit rate showed a significant difference (Initial task > Reverse task, p=0.009, t test). **c.** Each panel shows the stacked post-stimulus time histograms (PSTHs) for individual single-units. Red, blue and green indicate enhancement, suppression and no change in responsiveness, respectively. Responsiveness was defined as a change from spontaneous baseline activity during task performance. Vertical dotted lines show stimulus onsets. The left and right columns show the PSTHs for the Initial and Reverse task, respectively. The identity of the stimulus corresponding to each panel of PSTHs is shown to the right of each panel. **d.** The top and middle panels show the single-unit (gray) and population average (black) responses during the decision time-window, for the low and high tones, respectively, before and after reversals. During the task, the low tone reversed behavioral meaning from target (Tar) to non-target (NonTar), and vice versa for the high tone. Stars indicate significant differences. See main text for p-values. The bottom panel shows the same comparison for the noise, which was a non-target in both the Initial and Reverse tasks. Noise responses were not significantly different between tasks.

The decrease in hit rate was both statistically significant and characteristic of performance during the Reverse task, occurring in 12/14 recording sessions. These behavioral data indicate that impaired target recognition after reversals did not arise from differences in motivation to lick, since safe rates were high for both the Initial and Reverse tasks. Furthermore, separating the behavioral responses according to tone frequency sub-bands did not result in any significant dependence of task performance on the tone frequency (Figure 3c, bottom panel). It is important to note that between the Initial and Reverse task, the animals received a Priming block (see Materials and Methods), in which each trial consisted of only the low tone and noise, paired with freely available water, until the false alarm rate to the low tone fell below ~25%. In contrast, the high tone was not heard for a time period equal to the length of the Priming block (~15 minutes). The presence of this Priming block may explain why the false alarm rate did not significantly increase during the Reverse task, and instead, the hit rate decreased: the animals had many more rewarded trials in the Priming block just before the Reverse task to learn the low tone’s reversed behavioral meaning (non-target), whereas the high tone’s reversed behavioral meaning (target) occurred suddenly at the onset of the Reverse task. The effectiveness of priming in enabling reversal learning is evidenced by a significant decrease in no-go behavior to non-target low tones during the Priming block, compared to the target low tone no-go behavior in the immediately preceding Initial task (−60.4% ± 3.2%, p<0.001, t-test).

**Figure 3.**
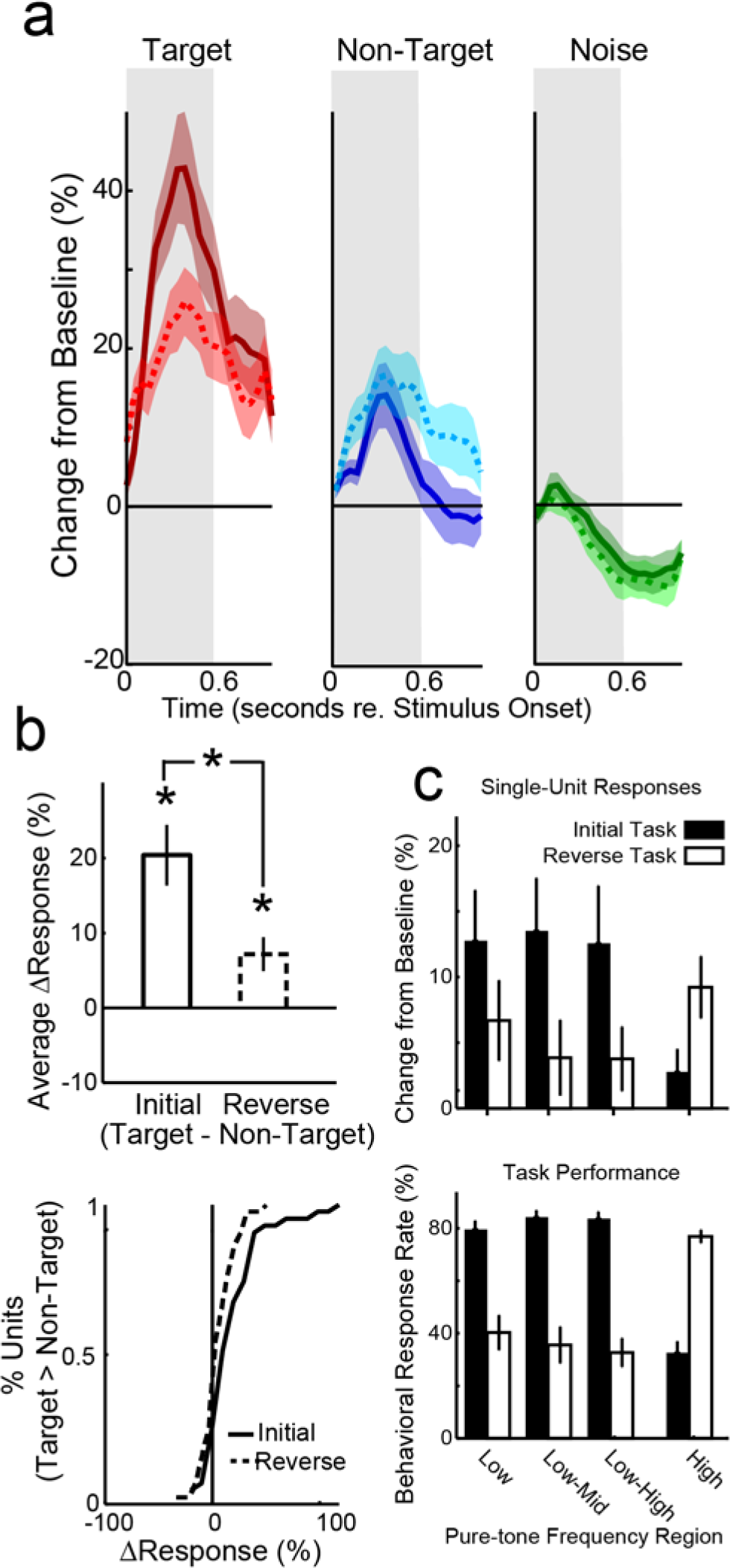
General linear model (GLM) of the dlFC single-unit population. **a.** GLM estimations of the population response time-courses for both the Initial and Reverse tasks are shown in solid and dotted lines, respectively. Target, non-target and noise responses are shown in red, blue and green, respectively. Shading shows the standard error of the mean (SEM). The gray regions show the 600 ms decision time-window within a trial. **b.** The top panel shows the difference between neural responsiveness (ΔResponse) during the behavioral decision time-window, for targets vs. non-targets in the Initial and Reverse tasks. Error bars show the SEM, and stars indicate statistically significant differences(bootstrapped t test—see Main text for p-values).The bottom panel shows the individual single-unit cumulative distribution function (CDF) for ΔResponse. **c.** Single-unit and behavioral data (top and bottom panels, respectively) were divided into tone frequency sub-bands. During the Initial task (black bars) the low and high tones were targets and non-targets, respectively. During the Reverse task (white bars) the high tones became targets, while the low tones became non-targets. Using the GLM to estimate neural and behavioral responsiveness as a function of tone frequency within target and non-target categories, we did not see any significant dependencies on tone frequency (p>0.05, bootstrapped t test).

Thus, a plausible explanation for the impaired target recognition after reversals is that the association of high frequency tones and approach behavior during the Initial task, interfered with the association of high frequency tones and avoidance behavior during the Reverse task. In other words, impaired target recognition during the Reverse task occurred because of proactive interference arising from a persistent sensory-cognitive association from the Initial task.

### Single-units adaptively represent behavioral meaning across reversals

During performance of the auditory reversal task, the majority of auditory units (68%) showed a common pattern of responses, as seen in the heat-map of post-stimulus time histograms (PSTHs) in Figure 2c. We observed strongly enhanced target tone responses, suppressed noise responses, and moderately enhanced non-target tone responses. This pattern occurred in both the Initial and Reverse tasks. We quantified the plasticity of each unit’s responses to each sound, by comparing the average percent change from baseline during the decision time-window, before and after reversals (Figure 2d). We found that when the low frequency target tones from the Initial task became non-targets during the Reverse task, neural responses significantly decreased (−10.4% ± 3.9%; p=0.033, bootstrapped t-test). The opposite effect occurred for high frequency tones: responses significantly increased after reversals (16.6% ± 4.5%; p<.001, bootstrapped t-test). In addition, we found that the animals’ decreased no-go behavior in response to low tones during Priming (i.e. evidence of reversal learning) was mirrored in dlFC by reduced neural responses to low tones during Priming (−8.3% ± 2.8%, p= 0.058, bootstrapped t-test). These data indicate that dlFC can arbitrarily map stimulus features to new behavioral meanings, which may facilitate reversal learning.

It is noteworthy that in contrast to the tone responses, the noise, which was an unambiguous approach signal, elicited a suppressive response in both the Initial task and Reverse task (blue in bottom panels of figure 2c—and see Figure 3a, explained below), but was not significantly different between the Initial and Reverse tasks (−8.5% ± 4.5%; p=.164, bootstrapped t-test). This may be the result of the constant neutral value of the noise stimuli across all task and priming conditions.

To compare the influence of stimulus frequency, stimulus behavioral meaning, and the animal’s behavioral decisions in determining the modulation of single-unit responses, we used a general linear model (GLM) of spiking for each single-unit. Figure 3a shows the GLM estimation of the dlFC population response time-courses, for target and non-target tones, and noise. The population averaged time-courses matched individual single-unit PSTHs in Figure 2c: strongly enhanced target tone responses, suppressed noise responses, and moderately enhanced non-target tone responses. Figure 3b shows the individual single-unit cumulative distribution functions (CDFs) and population averaged difference in dlFC response amplitudes (ΔResponse) for targets vs. non-targets. The population averaged target enhancement was significant in both the Initial task (20.4% ± 4.1%; p=0.0002, bootstrapped t-test) and Reverse task (7.2% ± 2.3%; p=0.049, bootstrapped t-test). However, the Reverse task had significantly less target enhancement than the Initial Task (p=0.007, bootstrapped t- test). The single-unit CDF shows that this finding was also true for most individual single-units: the Reverse task was skewed to the left of the single-unit CDF for the Initial task.

To rule out the possibility that specific tone frequencies may have biased neural or behavioral responses, we subdivided the single-unit and behavioral data into sub-bands of tone frequencies. As in our behavioral results, separating the neural responses according to tone frequency sub-bands did not result in any significant dependence of neural responses on tone frequency within targets or non-targets (Figure 3c). However, on average across the population, low frequency targets during the Initial task had significantly greater responses than high frequency targets in the Reverse task (ΔResponse=9.5% ± 1.4%; p=0.04, bootstrapped t test). There was no significant difference in response amplitude between low and high frequency non-targets (ΔResponse=3.7% ± 2.3%; p=0.31, bootstrapped t test).

### Single-unit responses during proactive interference

We found that dlFC single-units encode a sound’s behavioral meaning, and had smaller target responses after reversals. We suggest that decreased target responses during the Reverse task reflect proactive interference from the previous target-tone association in the Initial task. Since proactive interference led to impaired target recognition in the Reverse task, we predicted that the diminished neural representation of behavioral meaning after reversals might be most pronounced during incorrect behavioral responses to targets in the Reverse task. To test this prediction, we used the GLM to quantify neural responses to tones during the decision time-window in both the Initial and Reverse tasks. We grouped neural responses according to behavioral response-types: target hits and misses, and non-target false alarms and correct rejections. Figure 4a shows the GLM estimation of response modulation from baseline for each condition.

**Figure 4.**
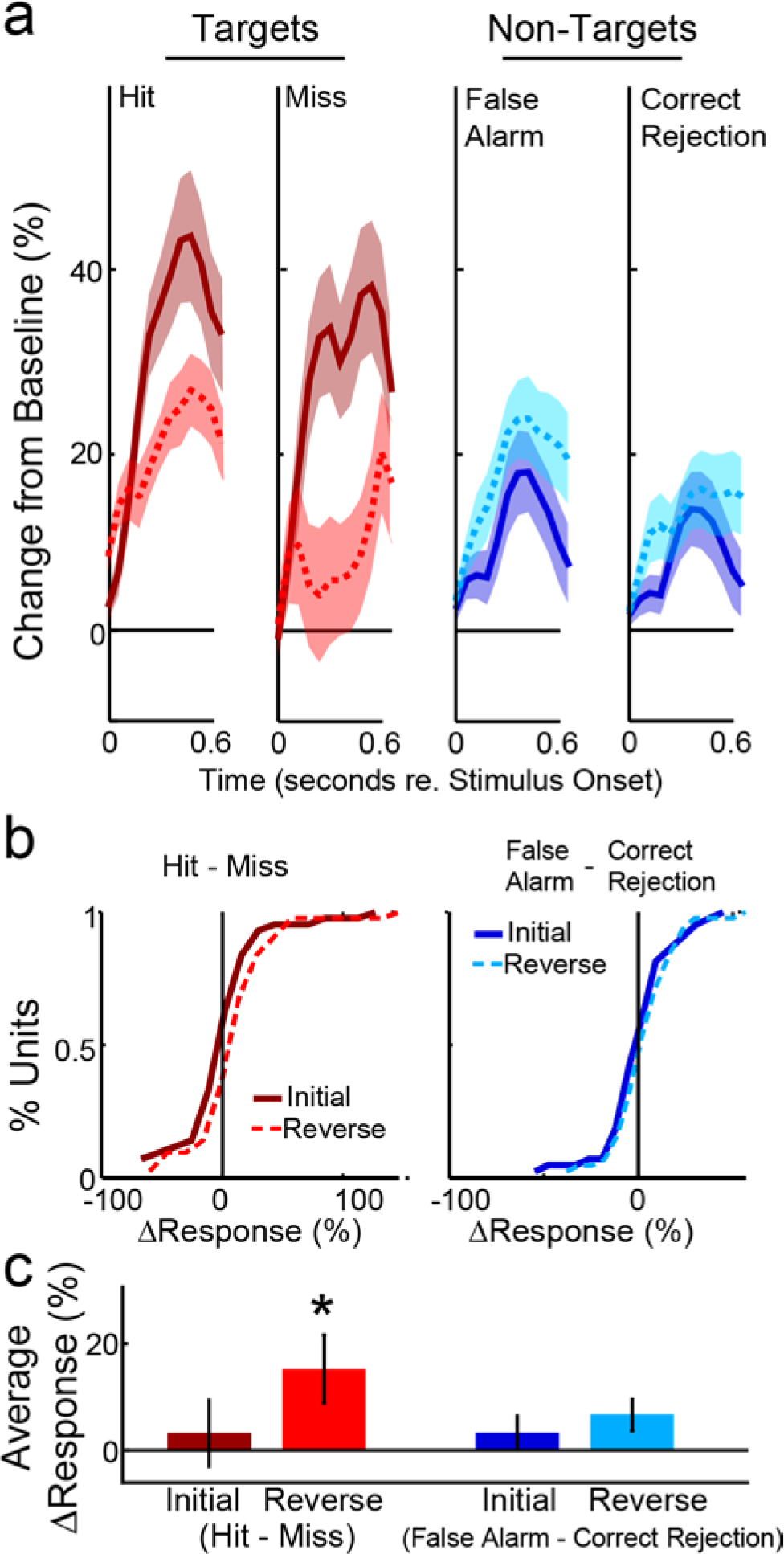
Single-unit adaptive plasticity in dlFC during proactive interference. **a.** Population results of the GLM evaluated for target and non-target tones (red and blue, respectively), during the Initial and Reverse tasks (dark solid and light dotted lines, respectively). Neural responses were grouped according to behavioral responses to tones: target hits and misses, non-target false alarms and correct rejections. Shading shows the SEM. **b.** Individual single-unit CDFs and **c.** Population average ΔResponse (here, hits vs. misses and false alarms vs. correct rejections), for the data color-coded as in Fig. 3A. Error bars show the SEM, and the star indicates the only significant difference (Reverse task target hits > target misses; p=0.04, bootstrapped t test).

As predicted, the smallest target responses occurred during target misses in the Reverse task. This can be seen in Figure 4b and c single-unit CDFs and population averages, where ΔResponse is plotted for hits vs. misses and false alarms vs. correct rejections. The hit vs. miss CDF in Figure 4b is clearly skewed to the right of 0 for the Reverse task, indicating that the majority of single-units were less active during incorrect behavioral responses to targets. The population averaged ΔResponse (Figure 4c) was also significantly greater during hits than misses (12.2% ± 5.1%; p=0.04, bootstrapped t test). Similar to the behavioral results, none of the other comparisons, i.e., hits vs. misses in the Initial task, and false alarms vs. correct rejections in both Initial and Reverse tasks, yielded significant differences. In other words, both the behavioral and neural deficits after reversals were specific to target responses.

## Discussion

We trained two ferrets to perform an auditory reversal task, while we recorded single-unit spiking from dlFC. To our knowledge, this is the first study to use an auditory reversal task in the head-fixed non-primate. Our success with this task was enabled by the novel use of a Priming block between the Initial and Reverse tasks. We found that both before and after reversals, animals were highly motivated to perform the task, and adjusted to the change in tone-reward associations after reversals. Nevertheless, the animals’ behavior indicated impaired target recognition after reversals, reflecting proactive interference (see hit rates in Figure 2b). In the dlFC, we found that units with responses time-locked to auditory events were selective for behavioral meaning (Figures 2 and 3). Although some earlier studies have emphasized the role of orbital frontal cortex (OFC) in reversal learning (Chang, 2014; Wilson et al., 2014), others have also noted activation of dorsolateral PFC during proactive interference in humans (Hyafil et al., 2009; Wolf et al, 2010) and in visual reversal tasks in monkeys (Asaad et al., 1998).

As in our behavioral results, ferret dlFC encoding of behavioral meaning was less robust after reversals (Figure 3), and diminished even further during incorrect decisions about the behavioral meaning of targets (Figure 4). Furthermore, we did not find a dependence of either neural or behavioral responses on any given frequency *within* the ranges of target and non-target tones (Figure 3c).

### Encoding of behavioral meaning in PFC responses

Previous single-unit studies in the PFC of monkeys and rodents trained on auditory (Karlsson et al., 2012) and visual reversal tasks (Asaad et al., 1998; Asaad et al., 2000; Donahue and Lee, 2015; Watanabe, 1992) have observed similar response properties to what we observed in the ferret, suggesting that the presence of neurons encoding behavioral meaning in PFC is conserved across mammalian species. In monkeys, Watanabe (1992) observed many classes of units that responded preferentially to different combinations of sensory, motor and cognitive attributes of the task. The units we found in the ferret dlFC that encode the behavioral meaning of sounds correspond best to Watanabe’s “auditory meaning” units, which responded preferentially to whether or not a stimulus was associated with a reward, and also reversibly encoded sensory-cognitive associations for high and low frequency tones. Also in monkeys, Asaad et al (1998) found that single-unit activity in PFC encoded the animal’s incorrect choice after reversals—a finding similar to ours, and others (Kim and Shadlen, 1999; Lee et al., 2009; Plakke et al., 2013; Rigotti et al., 2013; Rodgers and DeWeese, 2014; Russ et al., 2008). Here, our signal detection theory-based analysis of dlFC spiking has revealed a trial-averaged effect on how dlFC spiking is related to proactive interference during a reversal task. Future studies may provide further detail by using machine learning methods, such as support vector machines, in order to determine if dlFC spiking predicts trial-by-trial errors after reversals.

### The role of behavioral certainty in modulating PFC responsiveness

Despite the spectral overlap between the noise and tones in our reversal task, neural responses to both target and non-target tones increased from baseline (albeit, targets significantly more than non-targets), while responses to noise decreased from baseline. Since dlFC did not appear to encode auditory frequency-dependent features, but rather, behavioral meaning, *why then was the noise-response different than the non-target tone response?* The answer may be related to the animals’ degree of certainty about the behavioral meaning of these stimuli. During behavior, we found that the animals responded correctly to >99% of noise presentations, both before and after reversals—indicating a high degree of certainty about the behavioral meaning of noise. In contrast, tones were a more ambiguous signal, possibly representing “go” or “no-go” on any given trial. Kusunoki et al (2010) studied PFC responses to visual stimuli with “neutral” behavioral meanings (termed “consistent non-target” in their study), which are analogous to the noise stimuli in our task, since their neutral stimulus had a fixed behavioral meaning (always “safe” go-stimuli). While Kusunoki and colleagues did not find suppressed responses to neutral images, they did observe that the magnitude of PFC responses was graded according the likelihood of the image being a target, which was 0% for neutral images. They also observed intermediate responses to “inconsistent non-targets” (i.e. stimuli that had been experienced previously as a target), which is similar to the intermediate responses between noise and target that we observed in both our Initial and Reverse tasks. Thus, neural responses during our reversal task may have been shaped by the animal’s level of certainty about the behavioral meaning of tones and noise, gained from the training history. To test this hypotheses, future versions of our reversal task could include an third target tone frequency-band, with a fixed behavioral meaning (“no-go”), to provide a valence-balanced control for the fixed behavioral meaning of noise (“go”).

### Comparison between current and previous studies in ferret frontal cortex

In our previous study of neural responses in ferret frontal cortex during an auditory behavioral task (Fritz et al., 2010), recordings were made both in dlFC and in the premotor cortex (rostral anterior sigmoid gyrus, rASG). Recordings from the two areas were pooled because of similar responses. In a recent study of responses in ferret frontal cortex (Zhou et al., 2016) recordings were made in rASG during a visual discrimination task. In the present study, recordings were almost exclusively made in the dlFC, which is likely to account for the lower percentage of motor (lick) related neurons found in our current study (4%), compared to Fritz et al. (2010; 33%). Our previous study also found that in different populations of frontal cortical neurons, target tone responses could be either enhanced or suppressed relative to baseline. Zhou et al. (2016) similarly found both excitatory and suppressive target responses during their visual task. In our earlier study we observed little or no response to inconsistent non-targets or to consistent non-targets (reference noise).

The divergence of our current results with previous neurophysiological findings in dlFC is likely due to multiple differences in task stimuli, structure and reward. For example, Zhou et al. (2010) used a two-alternative forced choice visual task, with positive reinforcement and no reversals of sensory-cognitive associations. In contrast, we used a conditioned avoidance go/no-go auditory reversal task. An important difference with Fritz et al. (2010) is that in the present study, behavioral sessions with punishment (shock) were preceded by positive reinforcement priming phases; there were no such priming phases in our earlier study. Moreover, in the two auditory discrimination tasks utilized in the previous study (Fritz et al., 2010), the category of high tones and low tones did not have distinct behavioral meanings, and tone frequencies in both tasks were fixed rather than variable within session. In addition, the shock window overlapped stimulus presentation in the present study, whereas it occurred with a delay following stimulus offset in our previous study.

In summary, we find that single-unit responses in dlFC represent the behavioral meaning of sensory events. However, during proactive interference, the robustness and accuracy of this representation decreases. Thus, the persistence of task-inappropriate responses in dlFC reflects a lack of cognitive flexibility that may contribute to proactive interference.

## Acknowledgments

We thank Dr. Pingbo Yin and Diego Elgueda for their assistance with surgeries. This research was supported in part by NIH NIDCD F32 DC013722, RO1 DC005779, RO1 DC007657, and T32 DC000046.

